# Identifying priority areas for research and conservation of the largetooth sawfish *Pristis pristis* in Colombia and Panama

**DOI:** 10.1101/2021.01.14.426658

**Authors:** Juliana López-Angarita, Juan Camilo Cubillos-M., Melany Villate-Moreno, Annissamyd Del Cid, Juan M Díaz, Richard Cooke, Alexander Tilley

**Affiliations:** Fundación Talking Oceans, Bogotá, Colombia; Biology II, Aquatic Ecology, Ludwig-Maximilians-Universität München, Planegg-Martinsried, Germany; Ecological Genomics Group, Institute of Biology and Environmental Sciences, University of Oldenburg, Oldenburg, Germany; Fundación MarViva, Clayton, Ciudad del Saber 145-A, Panama City, Panamá; Fundación MarViva, Bogotá, Colombia; Smithsonian Tropical Research Institute, Panama City, Panamá; Sistema Nacional de Investigadores, Ciudad del Saber, Panamá; WorldFish, Penang, Malaysia

**Keywords:** Pristidae, Biogeography, Shovelnose rays, Historical Ecology, Museum specimens, Archaeozoology, Fisheries, Interviews

## Abstract

Sawfishes are considered one of the most endangered families of fishes in the world. Their diadromous ecology and vulnerability to fishing nets have brought most populations to the brink of collapse. Conservation of the surviving populations is hindered by the paucity of historic and contemporary catch and observational records, and assessments of suitable coastal and riverine habitats. Colombia and Panama are two of 14 countries considered as a high priority for the development of species-specific national legal protection of the critically endangered largetooth sawfish (*Pristis pristis*). To construct a baseline for the temporal and spatial distribution of the largetooth sawfish in Colombia and Panama, we collected historical records from museum databases and from literature over the past century, analysed available small-scale fisheries landings databases, and conducted interviews with fish workers in 38 locations across both countries. We found 257 records of sawfish occurrences across both countries between 1896 and 2015, with 69% of the records before the year 2000. The declining trend in the frequency of observations was corroborated by fishers, who reported fewer sawfish catches over the last 20 years. Using kernel density estimation of recent encounter locations, we identify potential hotspots that may represent extant populations of sawfish. These locations are broadly characterized by their remoteness and high mangrove forest cover. Given the length and cultural diversity of the Pacific coastlines of Colombia and Panama, and the inaccessibility of many of the communities, our findings provide important guidance to target rapid conservation and fisheries interventions to priority areas. We suggest that the relative success of community-managed fishing areas in the region makes this a valuable platform on which to build local stewardship of marine resources, while raising awareness of the need to safeguard critically endangered largetooth sawfish.

## 1. Introduction

All five extant species of sawfish are listed by the International Union for the Conservation of Nature (IUCN) as endangered or critically endangered with extinction (Dulvy et al. 2016), making them the most threatened family of marine fishes at time of writing (Dulvy et al. 2014, Braulik et al. 2020). Sawfishes, characterised by their tooth-studded rostra and large body size, are distributed globally in the tropics and subtropics inhabiting shallow coastal waters, such as estuaries, mangrove areas, and rivers (Harrison & Dulvy 2014). Sawfish amphidromy has been widely documented in rivers of Central America as well as natural (Lake Cocibolca, Nicaragua) and man-made lakes (Lake Bayano, Panama) (Thorson 1976, Richard Cooke 1994). Their widespread decline has been associated with their high catchability in fisheries given their easily entangling rostra (Simpfendorfer 2000), the extensive loss of their nursery habitats (Hossain et al. 2015, Haque et al. 2020), their high value in the fin trade market (McDavitt 2014), and their low intrinsic rates of population increase (Harrison & Dulvy 2014).

Considerable research has documented the vulnerable status of sawfish populations around the world, and their disappearance from many coastal areas. From Papua New Guinea (White et al. 2017) to Guinea-Bissau (Leeney & Poncelet 2015), the USA (Seitz & Poulakis 2006) and Bangladesh (Haque et al. 2020), among others. Despite the tendency of each generation to assume current biological states as the baseline, the long-term human impacts on marine species and ecosystems are widely recognized (Jackson et al. 2001). As such, there has been a call to incorporate historical data into assessments of change (McClenachan et al. 2012), and more recently, into conservation and management strategies (Engelhard et al. 2015). Historical data can be important to generate an understanding of the magnitude and rate of ongoing changes in the abundance of declining or locally extirpated species (McClenachan et al. 2012). Without historical data, it is not possible to state appropriate recovery targets for endangered species, understand the trajectories of decline, or reconstruct ecosystem baselines (Thurstan et al. 2015). In the case of sawfish, using bibliographic archival searches, and extinction analysis, Ferretti et al. (Ferretti et al. 2016) reconstructed the history of sawfishes in the Mediterranean Sea and confirmed that this family was present in the region despite previous research suggesting the opposite.

The historical and contemporary status of sawfish populations in Colombia and Panama is largely unknown, due to the lack of detailed information on fishery catches and observations, particularly for sharks and rays in the region. However, the Pacific coasts of Panama and Colombia present suitable ecological habitat for sawfish, which combined with the low human population density, may harbour remnant populations of *P. pristis*. A recent assessment of the global conservation status of sawfishes, (Dulvy et al. 2016) found that two species are present in Colombia and Panama. The globally distributed largetooth sawfish *Pristis pristis* (Linnaeus, 1758) is known to have occurred previously in the southern Caribbean (Santiago et al. 2014) and on the Pacific coasts of both countries. *P. pristis* is the largest species of the sawfish family, reaching a maximum length of 7.5 metres, and can live for up to 30 years (https://www.fishbase.se/). It is euryhaline, with adults preferring shallow marine and estuarine environments, and juveniles occurring in rivers (Thorson 1982). No recent biological or ecological information exists for *P. pristis* in Colombia and Panama, since it is an extremely rare species in the area, therefore difficult to assess and research. The species is known to be locally extinct in 27 of the 75 countries where it was historically present (Faria et al. 2013, Dulvy et al. 2016, Mendoza et al. 2017). In its native Caribbean range, *P. pristis* is thought to be locally extinct (Gómez-Rodríguez et al. 2014). In the Eastern Tropical Pacific, it could already be extinct along a large part of its original distribution from Mexico to Perú (Harrison & Dulvy 2014).

As of June 2019, the governments of Colombia and Panama agreed to impose strict national protections for all the sawfish species and cooperate regionally to recover populations through their classification as critically endangered under Annex II of the Specially Protected Areas and Wildlife (SPAW) Protocol. Colombia and Panama are two of the seven Caribbean countries categorised as the most urgent for the conservation of sawfishes (Harrison & Dulvy 2014, Koubrak 2016).

This study aimed to identify potential areas of extant populations of *P. pristis* in Panama and Colombia. By consolidating observation records from a variety of sources since records began, we identify historical changes in distribution, and by combining this with targeted interviews in key fisheries landing sites, we sought to identify the primary drivers of change. Given the combined Pacific coastline stretching more than 3000 km, this study is intended to form guidance to governments and stakeholder agencies in the region to identify starting locations to focus rapid conservation action, in support of their new commitments under the SPAW Protocol to foster the protection and recovery of populations of critically endangered sawfishes.

## 2. Materials and methods

### 2.1 Study area

Colombia and Panama are countries characterized by their high biodiversity and forest coverage, sharing regions of similar ecological and biogeographical features and special conservation value, such as the Chocó-Darien corridor. Both countries have both Pacific and Caribbean coastlines. Broadly, the Pacific coasts are characterized by rugged terrain, bays, swamps and sandy beaches. This area includes the Chocó-Darien ecoregion, considered to have one of the highest rates of annual rainfall in the world (c.a. 8000-13000 mm), feeding numerous river deltas and estuaries. In contrast with the Caribbean coast, the Pacific coast exhibits extensive mangrove habitat with four designated Ramsar wetlands i) Golfo de Montijo, ii) Bahia de Panama, iii) Punta Patiño (Darien) and iv) Baudo Delta (Chocó, Colombia) (https://www.ramsar.org/). This coastline is sparsely populated by Afro-Colombian and indigenous communities that rely heavily on small-scale and semi-industrial fisheries, subsistence hunting for wild game and agriculture for their livelihoods and food security.

Interview sampling was focused on the Pacific Coast of Colombia and Panama (Figure 1), as the types of ecosystems and habitats are favourable for largetooth sawfish, and previous research suggests they are currently locally extinct in the Colombian Caribbean (Gómez-Rodríguez et al. 2014, Caldas et al. 2017). Due to budgetary constraints, interviews were not conducted in the southern departments of Colombia lying on the Pacific coast (Valle del Cauca, Cauca and Nariño), but fisheries landings records were obtained and analysed from these areas.

**Figure 1.**
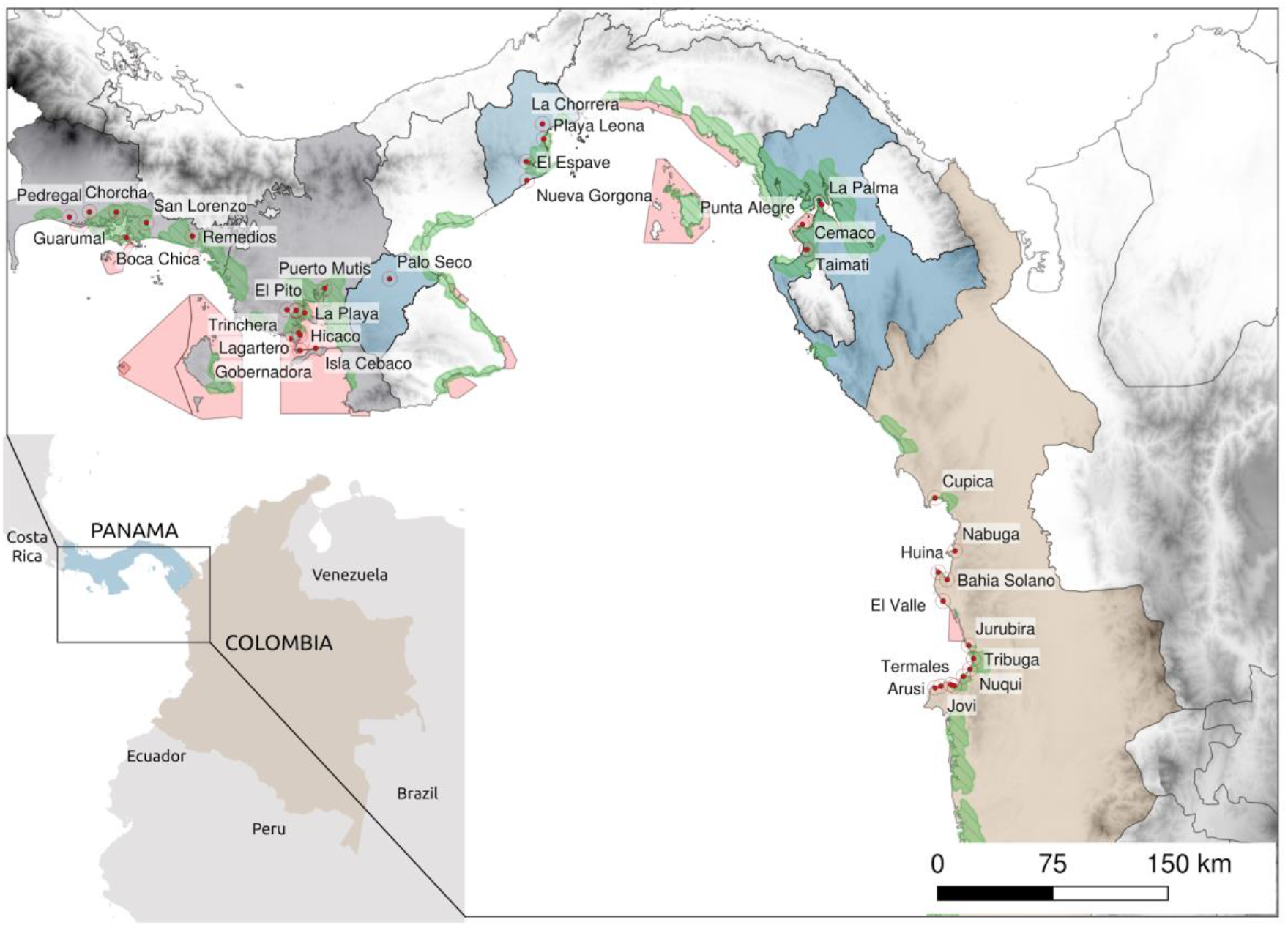
Study site map showing communities where interviews were undertaken between 2015 and 2016. Main regions surveyed in Panamá encompass the provinces of Chiriqui and Veraguas including the Bahia de Los Muertos and the Gulf of Montijo; and the provinces of Herrera, West Panamá, and Darién. For Colombia, interviews were carried out in the northern region of the Chocó department.

### 2.1 Historical sawfish records

A comprehensive literature review was undertaken using scientific and common names. Sawfish occurrences such as sightings, capture or other documented evidence of sawfish presence in Colombia and Panama were searched for in national and international online databases (Annex 1). Records found in the Global Biodiversity Information Facility (GBIF) were cross-referenced with museum collections to collect any additional information on capture location and date, and morphometrics of the specimens where available.

Relevant government departments and non-governmental organisations in both countries were visited or contacted to find recent and archaeozoological records documented in their databases (Annex 2). Museums in Colombia and Panama holding sawfish specimens were visited or contacted to collate information from collection metadata (Annex 1). Museum collection managers, curators and elasmobranch researchers were consulted directly to request information about possible specimens/ rostra, literature or expedition reports that could contribute more data, such as detailed capture location, year of capture, mode of capture, and specimen size and sex. Where preserved specimens existed and could be accessed, we measured morphometric variables such as total length (for full-body specimens), total rostrum length, and the number of teeth.

### 2.2 Fisheries catch records

Available fisheries landings data sets in Colombia and Panama were reviewed and explored for sawfish landings records. In Colombia, landings data in Zone C were collected by Fundación Marviva in a participative fisheries monitoring programme between 2010 and 2013 (López-Angarita et al. 2018a). For the southern Colombian Pacific coast, the landings database compiled as part of the USAID Bioredd+ fisheries program 2012-2014 (Tilley & Box 2014, Tilley et al., 2018) was consulted for sawfish records. In Panama, landings data were obtained through data-sharing agreements with Universidad de Panamá. These data were collected by the Panama government’s fishery resources authority ARAP (Autoridad de Recursos Pesqueros de Panamá), and are a record of fishery landings from 15 villages across the Gulf of Montijo between 2008 and 2012 (López-Angarita et al. in prep).

### 2.3 Interviews

The surveyed area encompassed the Pacific coast of Panamá and Colombia. In Panamá we surveyed communities across an area that stretches 772 km from the estuaries of Pedregal (Chiriqui, Panama) to Mariato in the estuaries of the Gulf of Montijo continuing to the Aguadulce District and finishing in Garachiné (Darien, Panama). In Colombia, we interviewed communities in the Northern chocó region between Punta Piña and Arusí. In total, we surveyed 38 communities (Figure 1).

Structured interviews were conducted by two trained scientists between 2015 and 2016. Local fish workers (i.e. fishers, traders, fishing cooperative representatives) and experts were asked to provide information on sawfish records or observations in or near their local area, according to their knowledge and experiences. Fishers with more than 15 years of fishing experience were considered to be local experts and were interviewed to collect fishers’ ecological knowledge data. Surveys were designed to gather information on: i) past and current abundance of sawfishes within the region, ii) perceived causes of decline, and vi) the cultural and economic importance of sawfish in their community. Photos of sawfishes were used to assess fish workers’ ability to identify the species, to record local names in use, and to prompt insights or records of catches and locations for the species. Printed maps were used to allow fish workers to identify specific areas of interest if they were too distant to show data collectors in person. Fish workers were asked to describe their encounters or sightings of sawfish. Where possible, date, locality, size and behaviour of the individual was recorded, however, if respondents struggled with specific details (for example from very old encounters), only specific data about their last sighting was used. Interviews were filmed with the informed consent of participants.

### 2.4 Data management and analysis

Occurrences of *Pristis microdon* and *Pristis perotteti* were treated as largetooth sawfish, *Pristis pristis* following the latest taxonomic reclassification (Faria et al. 2013). Any sawfish records from the Pacific coast were treated as *P. pristis*, whereas for the Caribbean coast, records were maintained as either *Pristis* sp., *P. pristis*, or *P. pectinata*. To clarify the process of sorting historical records, the term specimen is used to refer to records in museum collections. If these specimens could be associated with location and date information, then they are termed occurrences. Interviews were transcribed and coded by specific themes of interest. Qualitative analysis of interview responses was then performed using Nvivo software (version 11.1.1) to provide summary statistics of keywords and themes from the recorded answers.

Fisheries landings databases were filtered by species and location to provide a summary list of all sawfish captures along with capture date and location. The occurrences of sawfish in Panama and Colombia from bibliographic sources, museum specimens and interviews with associated coordinates were mapped using QGIS v. 3.8.0 (QGIS Development Team, 2019). Kernel density interpolation of geo-referenced occurrences was used to produce a heatmap of relative density (Heatmap v. 0.2). Interpolation was based on a kernel quartic (biweight) function with a cell size of 3 km and a decay ratio of zero. Each record was given an effective radius of influence (RD) of 50km (0.45 Degrees), heatmap coloration and gradient as the result of the cumulative effect of records in proximity lower to RD.

## 3. Results

A total of 252 records of sawfish encounters from 1896 to 2015 were compiled through museum specimens (59), the International Sawfish Encounter Database (24), literature (59), observations of interviewed fishers (95), and other non-interview observations (15) (Table 1). From these, 163 records belong to Panama (Pacific n = 149, Caribbean n = 3) and 88 to Colombia (Pacific n = 50, Caribbean n = 35). Twenty-one records had no accompanying date or location information. In Colombia, 65% of records date from before the year 2000, while this era represents the majority of occurrences recorded in Panama, with 82%.

**Table 1.**
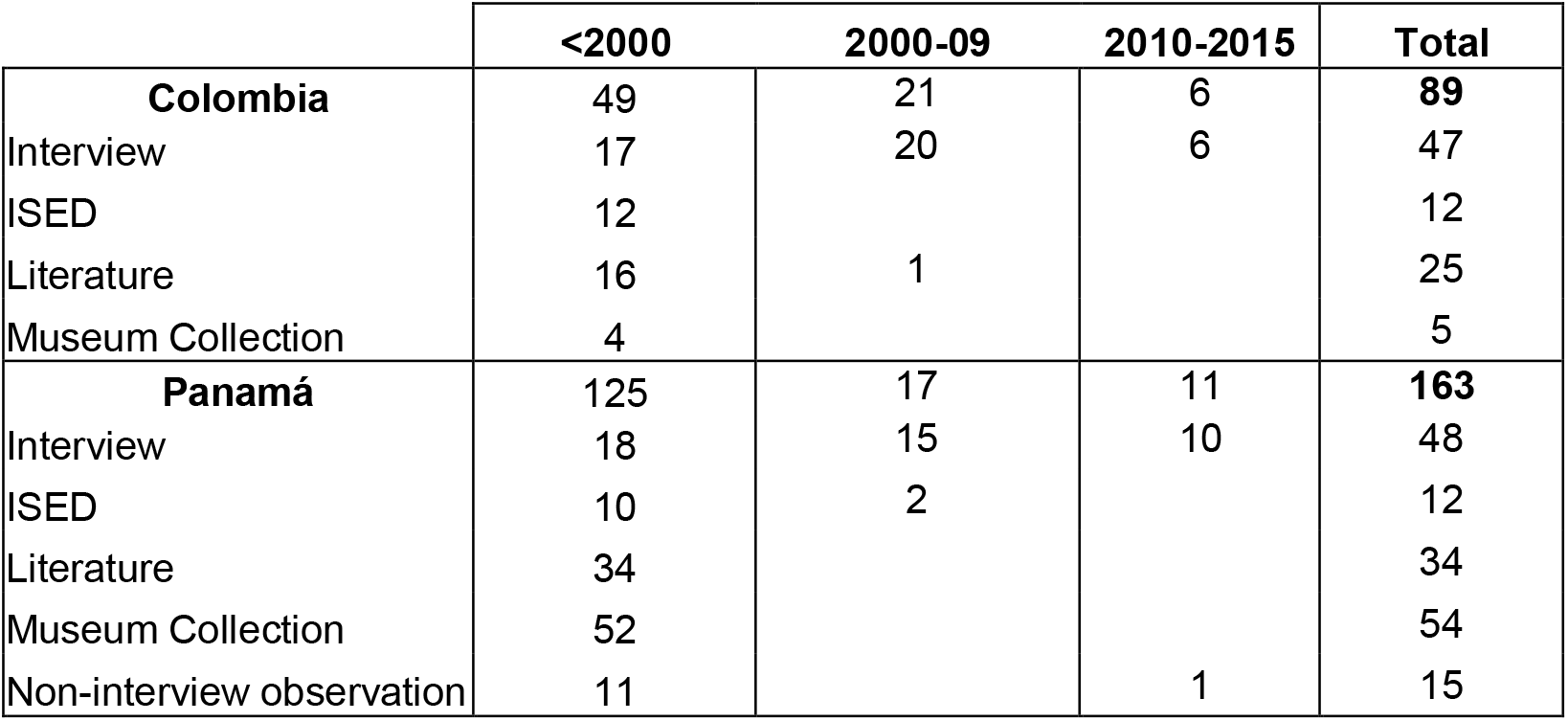
Counts of sawfish observations in Colombia and Panama (including Pacific and Caribbean coasts) according to record source type, location and year. ISED refers to records obtained from the International Sawfish Encounter Database, hosted by the Florida Museum of Natural History.

The total number of records with date and location data from the Pacific region across Colombia and Panama was 188. Of these, 71% correspond to observations before the year 2000, while 20% were reported between 2000 and 2009, and 9% since 2010 (Table 2). From the Caribbean region, 38 observations were obtained, from which 15 were largetooth sawfish (*P. pristis*), seven were smalltooth sawfish (*P. pectinata*), and the remaining were only catalogued to the genus *Pristis* sp. Largetooth sawfish, *P. pristis* is hereafter referred to as sawfish.

**Table 2.** Counts and proportion of largetooth sawfish (P. pristis) occurrences according to year for the Pacific region of Colombia and Panamá. Proportions exclude data with no associated date or location.

|  | <2000 |  | 2000-2009 |  | 2010-2015 |  | Total |
| --- | --- | --- | --- | --- | --- | --- | --- |
|  | Count | Proportion | Count | Proportion | Count | Proportion |  |
| **Colombia** | 20 | 42% | 21 | 45% | 6 | 13% | **47** |
| **Panamá** | 113 | 80% | 17 | 12% | 11 | 8% | **141** |
| **Total** | 133 | 71% | 37 | 20% | 17 | 9% | **188** |

### 3.1 Historical sawfish records (literature and collections)

Most specimens (wet and dry) found in collections and museums were housed outside Panama and Colombia, with most from the Field Museum of Natural History Fish Collection (Chicago, USA), the Smithsonian Institution’s National Museum of Natural History (NMNH) in Washington DC, and the Natural History Museum in London (UK) (Annex 1). The International Sawfish Encounter Database (ISED) yielded 24 records, of which all but one an accompanying date and location metadata. Of the 46 occurrences in the Pacific region, 44 belong to Panama, from which most are specimens from two biological expeditions (ichthyo-faunal assessments) in 1912 (Meek & Hildebrand 1923) and 1924. Four records were from Colombia, one was recorded as an observation in Guajira (Caribbean coast), and the other as a specimen in NMNH without a specified location. The other two records found in Colombia were found in the SIAM database for *P. pectinata* and *P. perotteti*, but no further information accompanied these records.

From 59 records found in bibliographic searches, four belong to archaeozoological evidence found in Colombia and Panama. The rest of the records were obtained from published literature dating back as far as 1890. For all of these, information regarding location, species and an approximate date were available. Of these records, 40 belong to *P. pristis*, seven to *P. pectinata* and 10 were only identified to genus. Morphological information was available for 30 records.

We contacted or personally visited 19 museums and zoological collections in the USA and Europe, seven in Colombia and three in Panama. Morphometric data were taken from six sawfish specimens found in Colombia and nine in Panama (Annex 1). Two of the specimens from Colombia did not have any accompanying information, but the whole specimen was available. Specimens from Panama lacked specific location data and consisted mainly of rostra. Information provided by the curators confirmed that specimens were brought to the museums before 2000.

### 3.2 Fisheries catch records

The database of small-scale fisheries landings from the northern Chocó, Colombia between 2010 and 2013 represented 36,448 fishing trips. The main gear types represented in the landings were: hand lines, long lines, gillnets, spear guns, beach seines and manual collection of molluscs and beach seine. Out of 449 reported fishing grounds (i.e. sites where people regularly fish), 32 were located inside mangroves, and 55 percent of the 284 species identified exhibit life histories that are strongly connected to mangrove habitat. However, no sawfish captures were reported. The three years of data available for small-scale fisheries in the Gulf of Montijo (Panamá) between 2010 and 2012 showed that sampling frequency and effort varied widely among the 14 villages sampled with large data gaps; some with entire years missing. Total catch volume across the three years was 203 tonnes and the most important fishery is “revoltura”, a local classification of the mixture of low-value species (e.g. weakfish, drums, croakers) caught with gill nets in nearshore mangrove and estuarine areas. No sawfish captures were reported.

### 3.3 Interviews

In total, 106 interviews were conducted across Colombia and Panama in 38 locations (Table 3). One hundred of the respondents were males and 6 were females. The mean number of years fishing (± SD) for all respondents was 36.5 years ± 12.5. The youngest respondent was 25 years old, and the oldest was 75 years.

**Table 3.** The number of interviews and communities by region and country conducted between 2015 and 2016. Location of the communities can be seen in Figure 1.

| Country | Region | Communities | Respondents |
| --- | --- | --- | --- |
| Panamá | Gulf of Chiriquí | 5 | 11 |
|  | Gulf of Montijo | 12 | 27 |
|  | Darien | 4 | 10 |
|  | West Panama City | 4 | 4 |
| Colombia | Northern Chocó | 13 | 54 |

Images of the largetooth sawfish were widely recognized by fish workers in Colombia and Panama, with 97% of respondents throughout the region correctly identifying the species. Local names for *P. pristis* in Colombia used by fish workers were: “pez sierra” (sawfish), “pez espada” (swordfish) and “guacapa”. In Panama, interviewees referred to the species as: “pez espada” and “pejapa”. 86% of respondents had seen a sawfish themselves, and even if they had not personally caught one, most knew someone who had. Sixty-six percent of respondents reported a decrease in the abundance of sawfish in their lifetimes. Twenty-eight percent of interviewees mentioned gillnets as possible reasons for sawfish decline, others suggested overfishing in general (16%), “extinction” (3%) and habitat degradation (2%) as additional reasons. Gillnets were highlighted as a particular threat because sawfish rostra become easily entangled, and some respondents recounted incidents of injury and fear while trying to extricate sawfish from their nets. In most locations, sawfishes had not been seen in more than 10 years (Figure 2).

**Figure 2.**
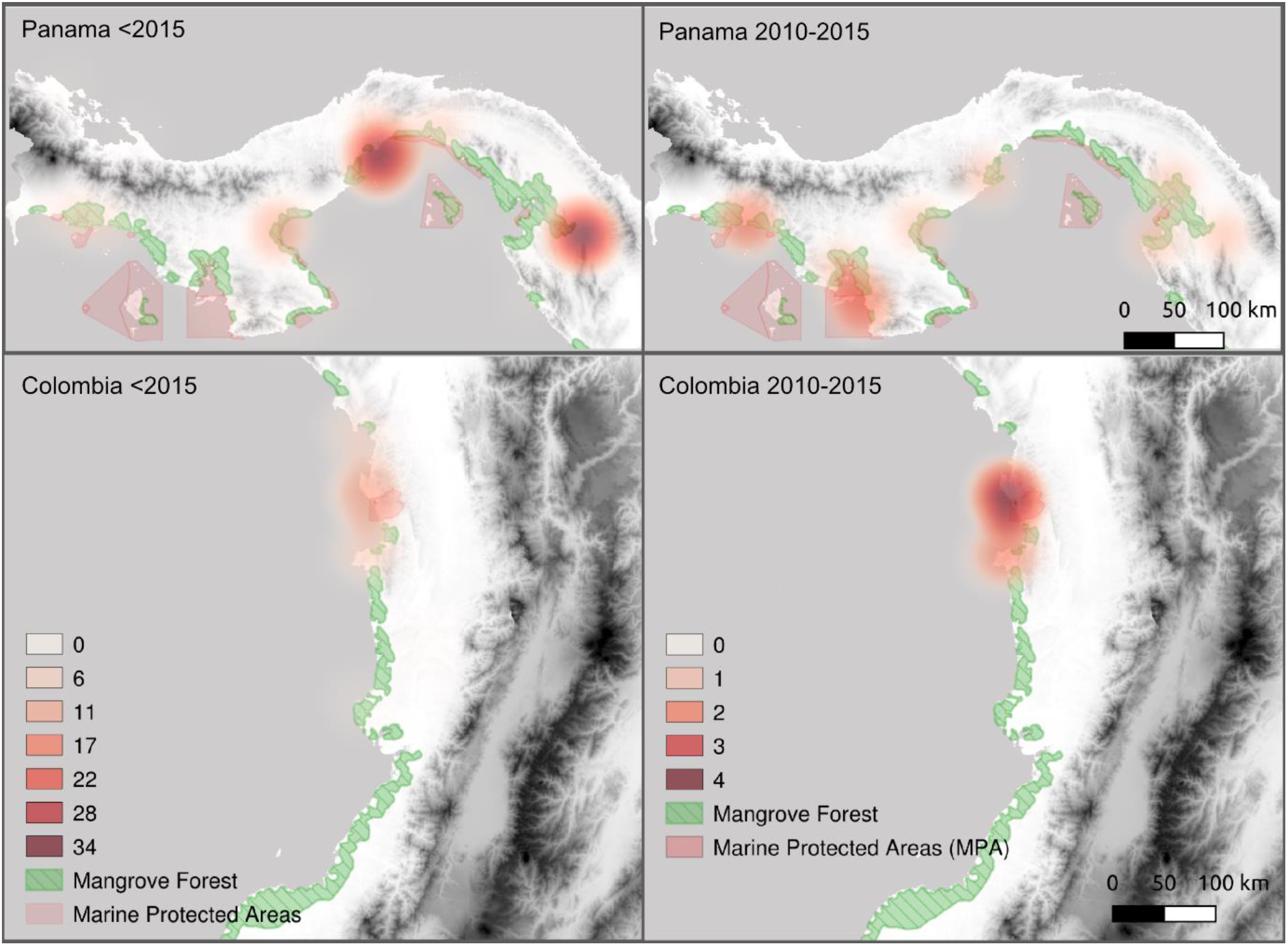
Heatmap (kernel density estimation) and distribution of largetooth sawfish (P. pristis) occurrences. All occurrences before 2015 for Panama and Colombia are shown in the left panel. The right panel depicts the distribution of sawfish occurrences only for years 2010 to 2015. Mangroves are highlighted in green and marine protected areas are shaded in red. Note the different gradient scales for occurrences numbers between panels.

Just over one-third of interviewees (38%) said they considered sawfishes to be very strong and dangerous animals that cause harm with their teeth or even attack and break fishers’ boats. Sawfish was never stated to be the target of any fisher and reported captures were on the whole perceived as accidental, opportunistic and even troublesome for fishers. Fish workers stated that they would usually keep the rostra of killed sawfishes as ornaments in their homes, but only one rostrum (45cm) was discovered during field sampling in the community of Punta Alegre (Darien, Panama), and fishers confirmed it was captured in 2015. Some interviewees claimed that in recent years many of the older rostra specimens had been purchased by visiting tourists or buyers. When asked their perceived reason for buyers’ interest in these rostra, 43% of interviewees stated it was merely decorative. For recent sawfish sightings, fisher workers mentioned two main locations for Panamá, the Darien region and the Bahía de Chame (La Chorrera región), and the Utria National Park and Golfo de Tribugá in Colombia.

### 3.4 Spatial-temporal analysis

A total of 150 occurrences from the Colombian and Panama Pacific region included a datestamp and location and were mapped for further spatial analysis. Occurrences from Panama were first reported at the beginning of the 20th century and are mostly composed of records collected during biological expeditions in the first part of the century. In Colombia, reporting only started in the 1980s, and most correspond to individual reports made by interviewees.

Occurrences of sawfish encounters in the last century are relatively rare and over 55% of the records are reported from three spatio-temporal clusters (Figure 2). Over a third of the occurrences (n = 60) belong to catches between 1912 and 1924 in Panama, collected in Panama Bay, West Panama, the Darien region and Taboga Islands. Another cluster in Panama represents 15 occurrences between 1985 and 2015 from Chiriqui (n=7), Cocle (n=4), Darien (n=2) and Veraguas (n=2), most of which correspond to single records and accounts from fishers. In Colombia, the largest cluster contained 11 occurrences recorded between 2000 and 2010 in the Utria National Park, the area with highest number of occurrences in the Northern Chocó (Figure 2). All occurrences here were reported to be adult sawfishes by interviewees.

## 4. DISCUSSION

In collating over a century of records of sawfish occurrences from Colombia and Panama, results show that sawfishes were historically distributed along the Pacific and Caribbean coastlines, but this distribution and their abundance appear to have decreased substantially with time. The paucity of recent observations and landings by fishers in most sampled areas, align with the widely reported global decline of sawfishes (Faria et al. 2013, Dulvy et al. 2016).

In both countries, the majority of records were dated from the 20th Century, with 79% of observations in Panama and 69% in Colombia occurring before the year 2000. The similarity between pre- and post-2000 records from the Pacific coast of Colombia can be explained by the few biological expeditions to this region. Interviewed fish workers stated that sawfish have never been commonly captured, but noted the species has been particularly scarce for the last 20 years. These results are consistent with recent research in the region, with the Colombian *Red List* reporting that *P. pristis* populations are possibly extinct from the Colombian Caribbean (Gómez-Rodríguez et al. 2014), and have been absent from the Pacific coast since the last recorded capture in 2007 (Caldas et al. 2017). However, our results suggest the Darien region in Panamá and the Northern Chocó region in Colombia are likely to retain local populations since they are hotspots of sawfish occurrences within the last 10 years. This is further supported by recent sightings reports. In November 2020, a ∼4m sawfish was observed swimming in the mangrove forest of Coquí in Northern Chocó (Ovidio Asprilla, *pers. comm*. nature guide from Coquí). Moreover, continuing captures of *P. pristis* in gill nets throughout 2020 have been reported at Rompío, a locality near a turbid marine inlet of the River Tuira in Darien (Panamá). Here, the sawfish is called *monã* in Emberá, the local indigenous language, and is consumed by local people (Nima Senapi, *pers. comm*. native Emberá from Unión, Darien).

More detailed spatial analysis shows that 40% come from expeditions in Panama between 1912 and 1924, specifically from Tuira River in Darien, Taboga Islands, Panama Bay and West Panama province. A third of these records belong to 19 young of the year (mean TL length 83 cm) and one neonate, while 12 were from adults (mean TL = 883 ± 106cm) (Meek & Hildebrand 1923, Breder 1927, Mitchell-Hedges 1928) when it was established that these areas were important sawfish pupping grounds.

Records reported for the last 5 years of our sampling period (2010 − 2015) make up only 13% of records in Colombia and 8% in Panama. The locations of these interviews overlap with occurrence hotspots from the spatial analysis, with fishers mentioning two main locations for Panamá; the Darien region and the Bahía de Chame (La Chorrera región), and the Utria National Park and Golfo de Tribugá in Colombia. The Utria National Park in the north of Colombia’s Chocó department was responsible for 6 reported records between the years 2000-2010. The northern Chocó is an area of particularly high biodiversity and it is divided into three different management areas; one national park and two community-managed fishing areas. These four sites are characterized by high coverage of mangrove forests and low human population density. Recent studies in the eastern tropical Pacific region have also documented sawfish sightings between 2013 and 2018 in areas of dense protected mangrove forest, such as the Térraba-Sierpe National Wetlands in Costa Rica (Valerio-Vargas & Espinoza 2019) and the Tumbes region in Northern Perú (Mendoza et al. 2017).

Sawfish records found in sites located 30 km from the coast in Malambo and 600 km upstream from the mouth of the Magdalena River (record from 1945, see Annex 1), illustrate how far upstream sawfish can travel. During interviews, fish workers mentioned that juvenile sawfish were generally found in river mouths and the shallow waters of small inland creeks, whereas adult sawfishes were caught during deep-sea industrial trawling. This suggests that large sawfishes are not restricted to shallow and coastal areas but maybe moving to deeper areas offshore that overlap with industrial fishery areas. Poulakis and Seitz (2004) found that the majority of adult sawfish encounters in deeper waters in Florida, occurred at the seabed level, indicating that deep trawling fisheries data are valuable for determining sawfish presence. This overall pattern of movement from shallow habitats towards open water areas away from the coast is consistent with data from encounter reports that have demonstrated a positive relationship between the increasing size of sawfish and an increasing mean depth/distance from the coast (Simpfendorfer and Wiley, 2005). This emphasises the need to obtain data on incidental catch from trawl fisheries for better understanding of threats and potential strategies to halt sawfish extirpation, but also that blame cannot be fully placed on coastal fishers, and that conservation efforts cannot be entirely localised.

The increase in the use of gillnets was the most frequent reason for sawfish decline highlighted by interviewees, while overfishing and habitat degradation, such as mangrove deforestation, were also mentioned. Fish workers stated during interviews that sawfish were not directly targeted, but rather were caught as by-catch in gill nets and generally died before being set free. In the Americas, coastal communities and mangroves have historically strong socio-ecological interactions, which have shaped the use of mangrove resources (e.g. shells, tannins, charcoal, fish) since PreColumbian times (López-Angarita et al. 2016). Human-sawfish interactions would have become suddenly more frequent with the emergence of gill nets as a common fishing gear in nearshore areas. Gill nets are pervasive in small-scale fisheries because they are cheap and effective, and continue to provide economically viable catch rates at dangerously high levels of effort (Northridge 1991). Elsewhere in the region gillnets have been identified as a major threat to sawfish (Chasqui et al. 2017, Mendoza et al. 2017), and in areas heavily fished with this gear, there is evidence of the historical reduction of sawfish populations, such as the Gulf of Nicoya in Costa Rica (Valerio-Vargas & Espinoza 2019).

Sawfish were not present in any of the artisanal fisheries data sets that we analysed for Panama (Gulf of Montijo) and Colombia (Northern Chocó, Southern Pacific coast). Interview findings indicate that sawfish were never abundant or captured frequently by fishers, hence the lack of records in fishery data sets may be a product of its intrinsic rarity, in addition to its widespread decline in the region. However, in other countries, largetooth sawfish are still persistent in fishery landings despite global decline trends, primarily as incidental catch of gillnets (Hossain et al. 2015, Haque et al. 2020).

For Colombia, localized fishery management schemes may play a role, as the database analysed for the Northern Chocó region includes data from a participatory monitoring program between 2010 and 2013, in an area that has restricted the use of nets since 2008 (Díaz & Galeano 2016). Gear regulations in this area have been regarded as effective in terms of compliance by fishers (Díaz & Caro, 2016) and fisheries stock recovery (López-Angarita et al. 2018a). Nowadays, the Pacific coast of Colombia has been peppered with management zones with similar gear regulations (Castellanos-Galindo & Zapata Padilla 2019). As sawfish mostly are found entangled in gillnets, the reduction in the number of recent catches seen in Colombia could be a positive effect of the reduced number of nets being used in this area.

The use of sawfish for meat, medicine, ornaments, among others, has been widely documented for several countries (McDavitt 2014, Leeney & Poncelet 2015, Leeney et al. 2018). Archaeozoological research in Panamá has shown sawfish were important for pre-Columbian cultures, not only as a source of food (Cooke & Jiménez 2008) but also for cultural and religious purposes, and thus, it possessed a significance that transcended their practical usage (Cooke 2004a b). Interviewees stated that once sawfish were caught, the meat was consumed and the rostrum was cut off and kept mainly for decorative purposes or later sold to buyers looking for their teeth. In Panama, some fishers reported that the teeth were fashioned into artificial spurs used in cockfighting, as documented from Perú (Cogorno Ventura 2001), something that has been observed by the authors on online sales sites and corroborated in other studies in Latin America (McDavitt 2014, Valerio-Vargas & Espinoza 2019). In Colombia, some fishers mentioned that the meat was used for consumption, but most pointed out that the rostrum was the most valuable sawfish body part because of its decorative value.

The eastern tropical Pacific has been highlighted in the agenda of sawfish global conservation because it has large, poorly studied regions with suitable sawfish habitat (Harrison & Dulvy 2014). Several fish workers stated that sawfishes were very frequent in the areas of Cuevita, Virudo and Pizarro in the southern Chocó region of Bajo Baudó, but it was not possible to survey these sites as part of this study. The coverage of potentially suitable habitat for sawfishes in this area is considerably wider than in the northern Chocó, with over 25,000 ha of estuaries and mangroves (Bernal et al. 2017). The area also includes the Baudo river delta, declared a Ramsar site in 2004 (Ramsar 2020). Historically, these areas were under substantial fishing pressure from industrial shrimp fisheries, suspended drift, and fixed gillnets, while lacking inclusive management zones for artisanal fishers (Velandia et al. 2019). However, with the prospect of reducing nearshore fishing pressure, community-based protected and management areas were established in 2017, in two mangrove areas (Frontera and El Encanto de los Manglares) (RUNAP 2018) resembling the community fisheries management implemented in Northern Chocó since 2008.

Future research efforts must focus on determining the status of any remnant sawfish populations in Bajo Baudo, especially since fisheries management legislation has been recently introduced. Another area of prime importance for future research is the Sanquianga National Park in the southern Pacific of Colombia; an expansive area of suitable habitat for sawfish with relatively low human density, that could also represent an undetected hotspot. In countries with historically high levels of mangrove deforestation such as Colombia and Panama (López-Angarita et al. 2016), management zones and protected areas have a crucial role in sawfish conservation, given that they not only may harbour the last functional populations of this critically endangered species, but they could guarantee its long term survival acting as a shield against habitat degradation (López-Angarita et al. 2018b) and non-selective fishing practices (i.e. gillnets) (López-Angarita et al. 2018a). Further research should leverage eDNA as a novel, non-invasive and rapid sampling technique to assess the presence of species (Simpfendorfer et al. 2016), especially in places where access and community surveys are restricted.

Remaining sawfish populations need critical management for sustained viability, yet without strong and urgent protection policy populations will continue to decline. Moreover, the implementation of the global strategy for sawfish conservation (Harrison & Dulvy 2014) will only be possible if interventions and regulations are established with local communities to ensure local legitimacy, enforcement and compliance (Rohe et al. 2017). A recent study from Bangladesh has illustrated this, where despite sawfish being protected under a national decree since 2012, they are still being landed at relatively high rates by fishers and traders unaware of the law (Haque et al. 2020).

There is widespread evidence that scientific research and published findings do not translate into effective conservation outcomes on-the-ground (Laurance et al. 2012). Actions must be put in place to overcome the gap between academic studies and their practical usability for conservation (O’Connell & White 2017). Therefore, in Colombia and Panama, building an understanding of the cultural and socio-economic value of sawfish with local communities in the context of their livelihoods and resource needs is the first step for their long-term conservation. Pushing sawfish onto national conservation agendas is a priority given the evidence that the species is still present in patches.

Our study supports other recent emerging research from the tropical eastern Pacific region in confirming the continuous distribution of the largetooth sawfish from Nicaragua to northern Perú (Mendoza et al. 2017, Valerio-Vargas & Espinoza 2019). The information compiled and generated here provides a strong baseline from which more detailed and exhaustive surveys can be undertaken. But more importantly for Panama and Colombia, it serves to put sawfish in the conservation agenda so that effective implementation of species protection protocols (i.e. SPAW) is achieved through working closely with fishers and local communities. This study highlighted several critical areas for sawfish recent occurrences, which is the first step in the process of creating evidence-based national conservation plans for this critically endangered species.

## 5. ACKNOWLEDGEMENTS

Firstly, we are indebted to all the local communities and respondents throughout Panama and Colombia for their patience and willingness to contribute their knowledge to this research. This work was carried out under a Keystone Grant from Save Our Seas Foundation (251). We extend our gratitude to the curators of natural history museums in Panama and Colombia for access to biological collections (Museo del Mar, Universidad Jorge Tadeo Lozano, Jose Julian Tavera from Universidad del Valle, Arturo Acero from Invemar). Thanks to Nestor Beltrán from the Humboldt Institute in Colombia for his help accessing databases. We greatly acknowledge Nicole Phillips (University of Southern Mississippi); Gavin Naylor and Tyler Bowling (Florida Program for Shark Research, Florida Museum of Natural History) along with Matthew McDavitt for all their valuable contributions to our historical record database. We also thank Caleb McMahan and Kevin Swagel (Field Museum) for their contributions, records and photographs in addition to all museums, collections, and NGOs that answered our data requests.

